# A novel strategy to mitigate *Corynebacterium bovis*-associated hyperkeratosis (CAH) in athymic nude mice

**DOI:** 10.1101/2025.05.15.654331

**Authors:** Abigail Michelson, Christopher Cheleuitte-Nieves, Kourtney Nickerson, Ileana C Miranda, Rodolfo J Ricart Arbona, Irina Dobtsis, Juliette Wipf, Neil S Lipman

## Abstract

Nude mice were inoculated with a non-pathogenic Cb isolate (NPI) or *Corynebacterium amycolatum* (Ca) to assess whether either could prevent skin lesions following inoculation with a pathogenic Cb isolate (PI). Crl:NU(NCr)-*Foxn1*^*nu*^ mice (n=6/group) were randomized into 6 groups: NPI (10^8^ CFU); Ca (10^8^ CFU); NPI or Ca followed 2 weeks later by PI (10^4^ CFU); and negative and positive controls receiving sterile media or the PI (10^4^ CFU), respectively. Colonization was assessed biweekly using isolate-specific PCR assays. Skin lesions were scored 0 – 5 daily for 4 or 6 weeks at which point skin biopsies were collected, evaluated and scored. No mice inoculated with the NPI and subsequently infected with the PI developed clinical signs nor was a significant amount of the PI detected by PCR. Mice inoculated with Ca before the PI developed milder, delayed skin lesions reaching a significantly lower mean peak clinical score (MPCS; 1.2 +/- 0.4) as compared to the positive control (MPCS 2.5 +/- 0.5). The Ca inoculated mice with and without PI had similar total histopathology scores, both of which were significantly higher than the mice inoculated with the NPI followed by the PI. These results led to evaluation of a practical exposure strategy in which nude mice (n=6/group) were housed on NPI-seeded bedding (SB) for 3 or 7 days prior to PI administration; mice housed on Cb-free bedding served as controls. Only 1 of 12 mice housed on SB receiving the PI developed CAH (peak score of 4), whereas all unvaccinated mice receiving the PI developed CAH (MPCS 2.83 +/- 0.69). The PI was not detected in the SB + PI groups until 21 days post-infection with the PI. There was no significant difference in total histopathology scores across groups, but the histopathology scores were lower in mice receiving the SB.

## Introduction

The gram-positive bacterial rod *Corynebacterium bovis* (Cb) causes Corynebacterium-associated hyperkeratosis (CAH), also known as scaly skin disease, in immunocompromised mice. This disease impacts the animal’s health as well as the research in which they are used.^1-3^ Cb infection results in 100% colonization with a variable clinical presentation depending on the isolate, dose, method of infection, as well as the age, strain, stock, and colony source of the mouse.^1,4-6^ Susceptible, immunocompromised strains and stocks demonstrate a diffuse scaling dermatitis and alopecia (in hirsute strains), and acanthosis and orthokeratotic hyperkeratosis microscopically with a relatively low mortality rate.^1,2,7^ Nude mice typically present with clinical signs 6-10 days post-infection which can include, in addition to the dermatopathology, weight loss, dehydration, erythema, and pruritus.^1,5,7,8^ Skin lesions typically resolve in nude mice 12 to 17 days post-infection, with some variation depending on dose, isolate, method of infection.^1,5,6,8,9^

Cb is logistically challenging to eradicate as it is often ubiquitous in vivaria housing immunocompromised mice.^2^ Eradication has been achieved using a depopulation and decontamination strategy, although these expensive, labor and time intensive efforts are often thwarted by the disease’s enzootic nature.^10,11^ Variable success in the management and eradication of Cb has been achieved using antibiotic therapy.^2,12-14^ However, once infected, immunocompromised mice remain colonized with Cb despite antibiotic treatment and in some cases, clinical resolution, and there can be consequences of long-term antibiotic use.^2,12,15,16^

Mendoza et al. recently demonstrated that not all Cb isolates evaluated were pathogenic to nude mice.^1^ Irrespective of an isolate’s pathogenicity, Cb isolates from mice and rats will colonize, but may not cause clinical CAH or significantly impact dermatopathology.^1,5,8^ They identified a Cb isolate (#24956) obtained from a vendor’s nude mouse production colony in which infected mice failed to develop clinically evident skin disease. It is unknown if the failure of these mice to develop clinically observable CAH or significant dermatopathology was a result of the isolate’s low pathogenicity, another unidentified factor, e.g., the skin microbiota or host genetics, or both.

A pilot study was subsequently undertaken to determine if administering this nonpathogenic isolate (NPI) to nude mice prior to exposure to a known pathogenic isolate (PI; #7894) would confer protection against clinical signs and/or histopathologic lesions associated with Cb infection. The PI Cb isolate had been shown to be the most pathogenic of all the Cb isolates in a study evaluating 9 isolates obtained from mice, rats, cows and humans.^1^ In the pilot study, the positive control mice who only received pathogenic Cb surprisingly did not develop clinical signs. The only difference between the pilot study and the study conducted by Mendoza et al. was the stock of outbred athymic nude mice utilized. Further investigation demonstrated that the stock and/or colony source can influence the presentation of CAH in outbred athymic nude mice.^5^ One trend that was found for mice that did not present with clinically detectable skin lesions was the presence of a known commensal of nude mouse skin, *Corynebacterium amycolatum* (Ca).

However, it remains to be determined if Ca alone, and/or in combination with other skin commensals or host genetics, influences the presentation of CAH.^5^ If a result of the skin microbiome, there are two theories as to how the microbiome provides protection. The first is based on competitive exclusion, by which two species relying on the same environment, in this case the skin, cannot stably or equally coexist.^17^ The second theory posits that the bacteria that provide protection prime the immune system, triggering the keratinocyte’s release of antimicrobial peptides, helping to control or resist infection with the pathogenic isolate.^18,19^ While the exact mechanism remains unknown, we hypothesized that the provision of either a NPI or Ca would confer protection against clinical disease associated with a PI Cb isolate.

## Materials and Methods

### Experimental design

#### Topical Inoculation

Nude (n=6/group; 3F, 3M) mice were randomly selected and divided into 6 groups (see Figure 1). Two groups were inoculated topically with 10^8^ colony forming units (CFUs) of either the NPI Cb #24956 or Ca #23001642. To evaluate the ability of these isolates to provide protection, 2 groups received either 10^8^ CFU NPI or 10^8^ CFU Ca and were subsequently challenged 2 weeks later with 10^4^ CFU of the PI Cb #7894. This PI dose was previously shown to cause disease in all inoculated nude mice.^1,5^ A negative control group received sterile culture media, and a positive control group received the same dose of the PI on the same day the NPI+PI and Ca+PI groups were inoculated with the PI. Colonization was confirmed via aerobic culture and PCR 7 days after inoculation with either NPI or Ca and every 14 days-post-inoculation (DPI) with the PI. The animals’ skin was visually assessed 5 times weekly and scored 0 – 5 based on lesion severity (Table 1) for 28 (Groups NPI, Ca, and sterile media) or 42 DPI (Groups NPI+PI, Ca+PI, and PI) at which point mice were euthanized by CO_2_, a gross necropsy was performed, macroscopic changes documented, and 4 full-thickness skin samples per mouse were collected, assessed histologically, and scored (Figure 4). An extended evaluation period of 42 days was utilized for the Cb+PI and Ca+PI groups to allow either the NPI or Ca to colonize prior to inoculation with the PI; the PI group was assessed following the same timeline.

**Table 1.** Cb clinical scoring for nude mice. Reprinted with permission from Mendoza et al.^1^

| Grade | Clinical signs |
| --- | --- |
| 0 | No clinical signs observed. |
| 1 | Mild amount of flakes/scales adhered to the skin and/or mild hunched posture with no erythema. |
| 2 | Mild amount of flakes/scales adhered to skin and/or mild hunched posture and mild erythema associated with flaking/scaly lesions. |
| 3 | Moderate erythema and/or moderate amount of white flakes adherent to the skin, or minimal white flakes on 3 or more additional locations affected including the ventrum, muzzle, cheeks, crown, and limbs. |
| 4 | Moderate erythema and/or moderate amount of white flakes adherent to the skin and moderate hunched posture. |
| 5 | Severe amount of flakes/scales adhered to the skin (thick layer), and/or severe erythema, and/or severe hunched posture. |

**Figure 1:**
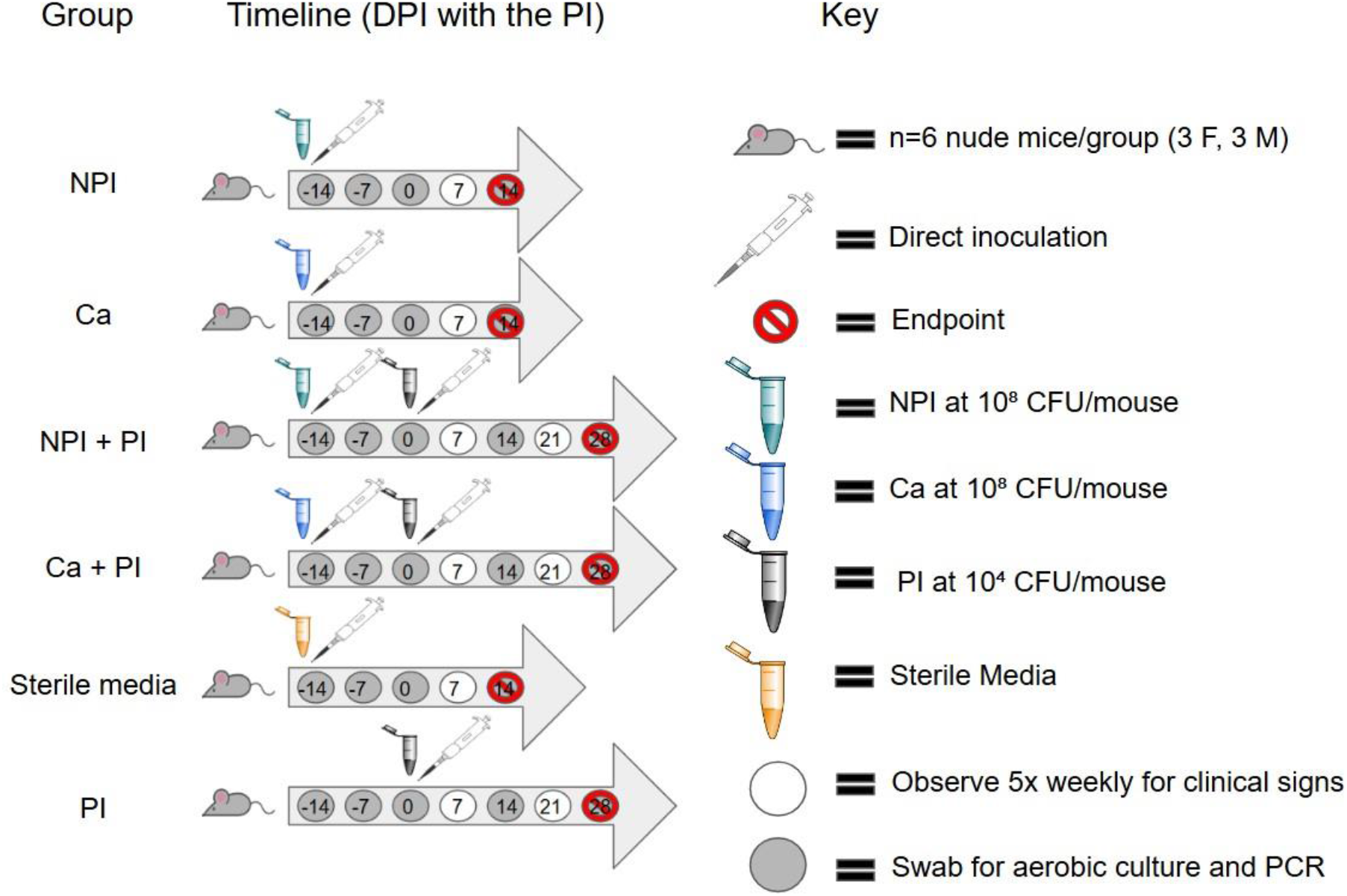
Schematic of topical inoculation study. Groups NPI and Ca received 10^8^ CFU/mouse of either the NPI or Ca. Groups NPI+PI and Ca+PI received NPI or Ca and were challenged 14 days later with 10^4^ CFU/mouse of the PI. The sterile media group received only sterile media, and the PI group received 10^4^ CFU/mouse of the PI at the same timepoint as groups NPI+PI and Ca+PI. DPI=days post-inoculation; NPI=nonpathogenic Cb; Ca=*Corynebacterium amycolatum*; PI=pathogenic Cb

#### Seeded Bedding Exposure

Nude mice (n=6 females/group) were divided into 4 groups (Figure 2). Groups SB7+PI and SB3+PI each received NPI seeded bedding (SB) and were subsequently inoculated with the PI (10^4^ CFUs/mouse) 7 (SB7 + PI) or 3 (SB3+PI) days after housing on the NPI SB. Group PI received sterile bedding and later exposed to the PI via direct inoculation at the same time and dose as the SB7+PI and SB3+PI groups. The SB group received the NPI SB at the same time as SB7+PI but was not inoculated with the PI. Groups SB7+PI, SB3+PI, and PI were all inoculated a second time with the PI (10^4^ CFUs/mouse) 35 days after the initial PI inoculum to evaluate the durability of protection provided by the NPI. Clinical signs, colonization, and histopathology were assessed for 70 days (63 DPI following the initial challenge) after housing on SB as described for the topical inoculation study.

**Figure 2:**
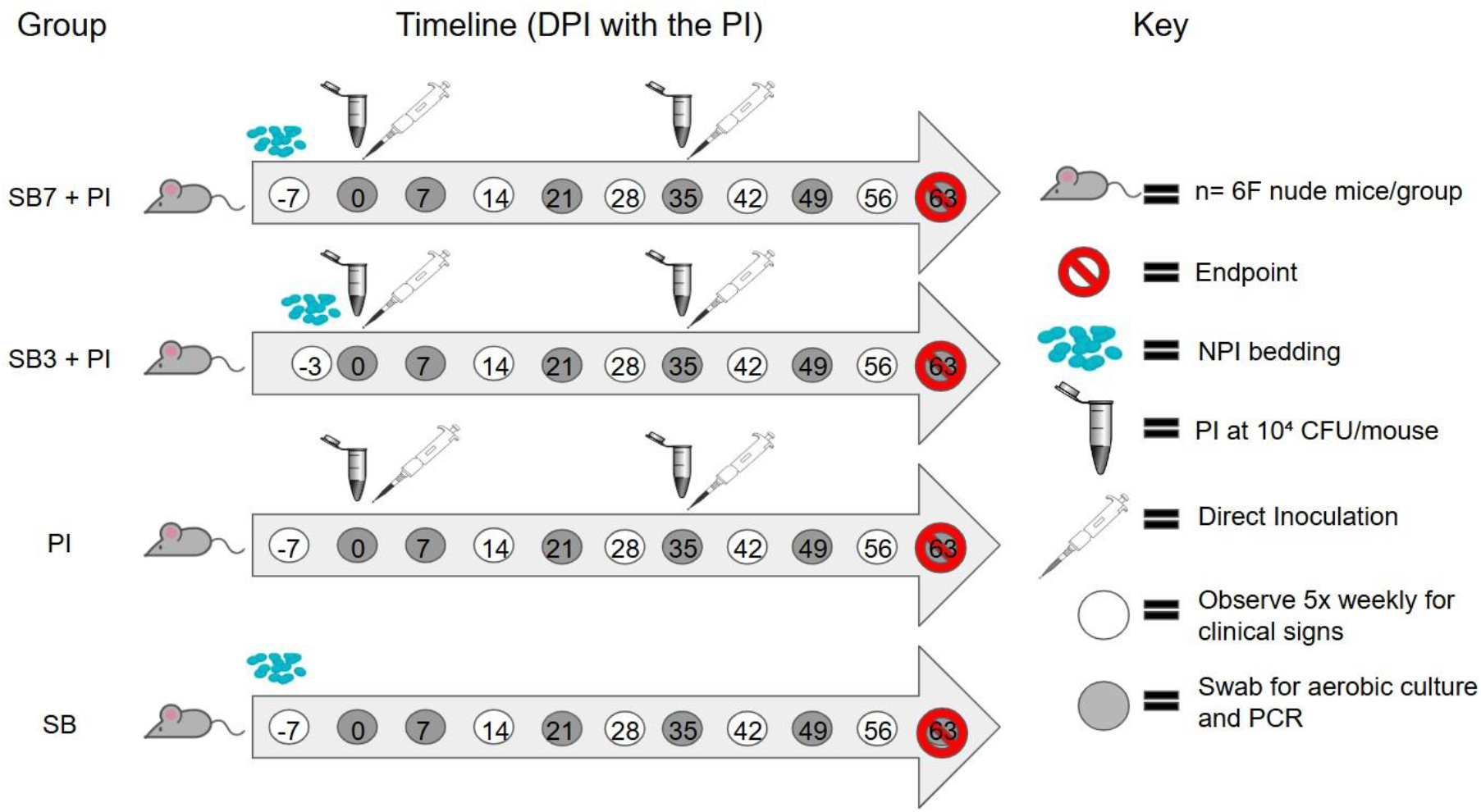
Schematic of seeded bedding exposure study. SB7 + PI group received the NPI seeded bedding 7 days prior to the initial inoculation with the PI. SB3 + PI group received NPI seeded bedding 3 days prior to the initial inoculation with the PI. PI group received sterile bedding while the SB group received the NPI seeded bedding at the same time as the SB7 + PI group but was not inoculated with the PI. Groups SB7+PI, SB3+PI, and PI were all inoculated with the PI a second time 35 days after the first PI inoculation.

### Animals

Forty-two female and 18 male, 6-to 8-week-old, outbred athymic nude mice (Crl:NU(NCr)-*Foxn1*^*nu*^, Charles River Laboratories, Wilmington, MA) were used. Only female mice were utilized for the seeded bedding exposure study due to the propensity of males to fight and the finding that there were no significant sex associated differences observed in the topical inoculation study. Mice were confirmed free of Cb infection upon arrival via aerobic culture and PCR. Mice were free of the following agents: *Respirovirus muris* (Sendai virus), *Orthopneumovirus muris* (PVM), Betacoronavirus muris (MHV), Protoparvovirus rodent1 (MPV/MVM), Cardiovirus theileri (TMEV), reovirus type 3, epizootic diarrhea of infant mice (mouse rotavirus), murine adenovirus, murine astrovirus-2, polyoma virus, K virus, murine cytomegalovirus, mouse thymic virus, lymphocytic choriomeningitis virus, Hantaan virus, ectromelia virus, lactate dehydrogenase elevating virus, Genogroup V norovirus GV (MNV), and Chaphamaparvovirus rodent1 (RCHPV-1), *Bordetella bronchiseptica, Citrobacter rodentium, Clostridium piliforme, Corynebacterium bovis, Corynebacterium kutscheri, Filobacterium rodentium, Mycoplasma pulmonis, Chlamydia muridarum, Salmonella spp*., *Streptobacillus moniliformis, Encephalitozoon cuniculi*, ectoparasites, endoparasites, and enteric protozoa.

### Husbandry and housing

Mice were housed in autoclaved, solid bottom and top, gasketed and sealed, polysulfone, positive-pressure, individually ventilated cages (Isocage, Allentown Caging Equipment Company, Allentown, NJ) on autoclaved aspen chip bedding (PWI Industries, Quebec, Ontario, Canada) in distinct cages based on group at a cage density of 3 mice per cage. Each cage was ventilated with HEPA-filtered air (filtration at rack and cage level) at approximately 30 air changes per hour and the HEPA-filtered rack effluent was exhausted directly into the building’s HVAC system. Mice received gamma-irradiated, autoclaved feed (5KA1, Purina Mills International, St Louis, MO) and autoclaved acidified (HCl) reverse-osmosis purified water (pH 2.5 to 2.8) ad libitum. Each cage was provided with a sterile bag of Glatfelter paper containing 6g of crinkled paper strips (EnviroPAK, WF Fisher and Son, Branchburg, NJ) for enrichment. Cages were changed every 2 weeks, and all manipulations were performed in a class II, type A2 biologic safety cabinet (BSC; LabGard S602-500, Nuaire, Plymouth, MN) using aseptic technique with sterile gloves. Media inoculated control mice were handled before experimental animals. Mono-inoculated animals were handled prior to handling mice inoculated with 2 bacterial species; mice inoculated with the NPI were handled prior to mice inoculated with the PI. The animal holding room was ventilated with 100% fresh air at a minimum of 10 air changes hourly and maintained at 72±2°F (21.5±1°C), relative humidity between 30% and 70%, and a 12:12-h light: dark photoperiod (lights on at 0600, off at 1800). All animal use was approved by Memorial Sloan Kettering’s (MSK’s) IACUC and conducted in accordance with AALAS’s position statements on the Humane Care and Use of Laboratory Animals and Alleviating Pain and Distress in Laboratory Animals with clear clinical endpoints described in the approved animal use protocol. MSK’s animal care and use program is AAALAC-accredited and operates in accordance with the recommendations provided in the Guide for the Care and Use of Laboratory Animals (8^th^ edition).^20^

### *Corynebacterium bovis and amycolatum* propagation and inoculation

Cb isolates #25496 (NPI) and #7894 (PI), and Ca #23001642 stored as frozen stock were grown on trypticase soy agar supplemented with 5% sheep blood (BBL TSA II 5% SB, Becton Dickinson, Sparks, MD) at 37°C with 5% CO_2_ for 48h, as previously described.^1,5,21^ The NPI and Ca were isolated from clinically normal nude mice from a vendor’s nude mouse production colony; the PI was isolated from a nude mouse with CAH at MSK. The growth curve for the Cb isolates had been previously established and the same method was used to determine the growth curve of Ca.^1^ The concentration of bacterial suspensions was approximated by comparing the culture’s absorbance using a MacFarland densitometer to a standard curve generated by serial dilutions.^1^ Bacterial suspensions used as inocula were collected from the cultures during mid-log of the growth phase, titrated and serially diluted in brain heart infusion broth (BHI; Becton Dickinson) supplemented with 0.1% Tween 80 (VWR Chemicals, Solon, OH) to obtain 10^8^ (NPI or Ca) or 10^4^ (PI) +/- 15% bacteria/50µl as previously described.^1^ For both the topical inoculation and seeded bedding studies, the challenge dose of the PI was based on findings from Mendoza et al.^1^

#### Topical inoculation

The bacterial inocula were placed in sterile 2ml polypropylene centrifuge tubes (Fisher Scientific, Waltham, MA), transferred to the vivarium on ice, and the animals were inoculated within an hour of inoculum preparation. Inoculation was performed in a class II, type A2 BSC (LabGard S602-500, Nuaire) using sterile materials and aseptic methods by personnel donning sterile gloves. After inoculation, the concentration of each inoculum was reconfirmed and enumerated by plate count. For topical inoculation and subsequent sample collection for aerobic bacterial culture, media inoculated control mice were handled before experimental animals. Each cage containing an experimentally naïve animal was removed from the rack and all sides of the cage were liberally sprayed with disinfectant (Peroxigard [1:16], Virox Technologies Inc., Oakville, ON) and placed in the BSC. Each cage was opened for less than 2 minutes. Bacterial inoculation was performed with the mouse restrained with the handler’s non-dominant hand by gently grasping the tail base and the animal’s hindquarters were elevated slightly while the animal grasped the wire bar lid with its forelimbs. The lid was positioned to allow the restrained animal to remain in the horizontal plane. Using the dominant hand, the bacterial inoculum was applied directly to the skin using a sterile filter micropipette (P200N, Marshal Scientific, Hampton, NH) along the dorsal midline starting at the nuchal crest proceeding to the tail, a distance of ∼ 2 cm. A 50μl bacterial suspension was evenly distributed along the dorsal midline at 4-5 sites dispensing 10-15μL per site using a sterile filtered 200uL pipette tip (Filtered Pipet Tips, Crystalgen, Commack, NY). This method ensured the bacterial suspension did not flow off the animal. The cage was closed and sprayed thoroughly with disinfectant prior to returning it to the rack. All interior surfaces of the BSC were sprayed with disinfectant and a new pair of sterile gloves were donned between each cage. A new cage was placed in the BSC as described and the steps repeated until all inoculations were completed. Animals in the control group were inoculated with an equivalent amount of bacterial-free BHI broth.

#### Seeded bedding

To generate NPI seeded bedding, the inoculum was grown to mid-log and diluted and titrated to 10^8^ CFU/ml. The inoculum (1.5ml) was mixed with sterile autoclaved bedding (45ml) in a 50ml sterile conical tube (Corning™ Falcon™, Fischer Scientific, Hampton, NH). The tube was vigorously shaken and then vortexed for 20 seconds to allow complete dispersal of the inoculum throughout the bedding. The conical tubes were stored in a -80°C freezer for 1 week before use. A pilot study confirmed that the bacterium could be recovered from the NPI seeded bedding after freezing at -80°C for 1 month. On the day of administration, the tubes containing the seeded bedding were thawed for at least 15 minutes. The tubes were shaken and vortexed as previously described and the bedding was administered to the mice within 1 hour of thawing. Each cage containing naïve animals was handled aseptically as previously described and 15mls of seeded bedding was added to each cage. Approximately 5 × 10^7^ CFU of the NPI were present in each 15 ml dose of seeded bedding. Each cage was opened for less than 2 min.

### Sample collection for culture and PCR

All mice were confirmed negative for Cb and Ca by aerobic culture and PCR utilizing isolate specific PCR assays prior to commencing the study. Mice were then sampled 14 days after inoculation with either NPI or Ca immediately before inoculating mice with the PI (0 DPI) and every 14 days for the duration of each study group (topical inoculation: 28 DPI with the PI; seeded bedding: 63 DPI with the PI). To prevent cross-contamination within each dose group, media inoculated controls followed by animals with lower clinical scores were swabbed first. When animals had similar scores, cages were selected at random. All cages were handled aseptically as previously described. Each cage was opened for no longer than 5 min. The animal was grasped gently by the base of the tail, allowing all 4 limbs to grasp the wire bar lid. A sterile culture swab (BBL CultureSwab, Becton Dickinson) was held perpendicular to the animal and swabbed along the right dorsal midline from the base of the tail cranially to the nuchal crest and caudally back to the tail base, rotating the swab as it was advanced. Samples for PCR were collected by rubbing an individually packaged adhesive swab (PurFlock ULTRA, Puritan Medical Products Company LLC, Guilford, ME) along the left side of midline, extending from the base of the tail cranially to the nuchal crest and caudally back to the tail base rotating the swab as it was advanced. Swab tips were broken into 1.5mL microfuge tubes and stored at -80°C prior to testing.

### Aerobic culture

Skin swabs were each streaked in a 4-quadrant pattern onto 3 agar plates: TSA with 5% sheep blood (TSA II 5% SB, Becton Dickinson), CNA with 5% sheep blood (Columbia CNA 5% SB, Becton Dickinson) and an additional TSA plate with 5% sheep blood overlaid with 1 drop of Tween 80.^22^ The plates were incubated for 72h at 37°C with 5% CO_2_. When present, distinct colonies were speciated by using MALDI-TOF spectroscopy (MALDI Biotyper Sirius CA System, Bruker, Billerica, MA). Plates without growth were held for an additional 7 days before being considered negative. Plates were scored from + to ++++ based on the number of quadrants with bacterial growth.

### PCR assays and targeted sequencing

Fluorogenic RT-PCR assays were developed targeting the specific Cb isolates utilized and Ca. To identify unique regions for PCR assay generation, genomic sequences from Cb isolates #24956 (NPI) and #7894 (PI) and Ca were aligned to publicly available sequences for Cb and closely related bacteria. Specific differences between isolates #24956 (NPI) and #7894 (PI) were identified as candidate regions for assay design to differentiate the strains. These regions were assessed to determine cross-reactivity for other Cb isolates and nucleic acid sequence by BLAST analysis against the nucleotide database or whole-genome sequence database gated on *Corynebacterium*. A total of 8 assays representing 4 targets for each strain were identified for development of proprietary real-time fluorogenic 5’ nuclease PCR assays. All 8 assays were qualified and optimized for use and after demonstrating similar performance, single assays for #24956 and #7894 were chosen. This process was repeated using target regions identified for Ca for development of a single assay. A commercially available, broadly targeting Cb PCR assay was used as a reference (Charles River Laboratories, Wilmington, MA). The assay was used to determine the presence of genomic DNA in samples as previously described.^23^ Samples that amplified during initial testing were subsequently retested using DNA isolated from a retained lysate sample to confirm the original finding. A positive result was reported when the retested sample was confirmed positive. To monitor for successful DNA recovery after extraction and to assess whether PCR inhibitors were present, an exogenous nucleic acid recovery control assay was added to each sample after the lysis step and prior to magnetic nucleic acid isolation. The concentration of eluted nucleic acid in mock extracted samples (no sample material) was calibrated to approximately 40 copies of exogenous DNA/µL and compared with a 100-copy system suitability control. A second real-time fluorogenic 5’ nuclease PCR assay was used to target the exogenous template to serve as a sample suitability control and was performed simultaneously with the Cb assay. Nucleic acid recovery control assays for samples that had greater than a log10 loss of template copies compared with control wells were diluted 1:4 and retested, reextracted or both prior to accepting results as valid. A 100-copy/reaction positive control plasmid template containing the Cb target template was co-PCR amplified with the test sample to demonstrate master mix and PCR amplification equipment function.

### Clinical scoring

Mice were assessed 5 times weekly throughout the duration of the studies. Animals were evaluated cage side and the mouse’s entire dorsal surface was assessed and scored using the system utilized by Mendoza et al. and provided in Table 1 and Figure 3.^13^ Mean peak clinical scores (MPCS) were computed by averaging all peak scores demonstrated by the mice in each respective group by day. The area under the curve (AUC) was also determined and compared per group.

**Figure 3:**
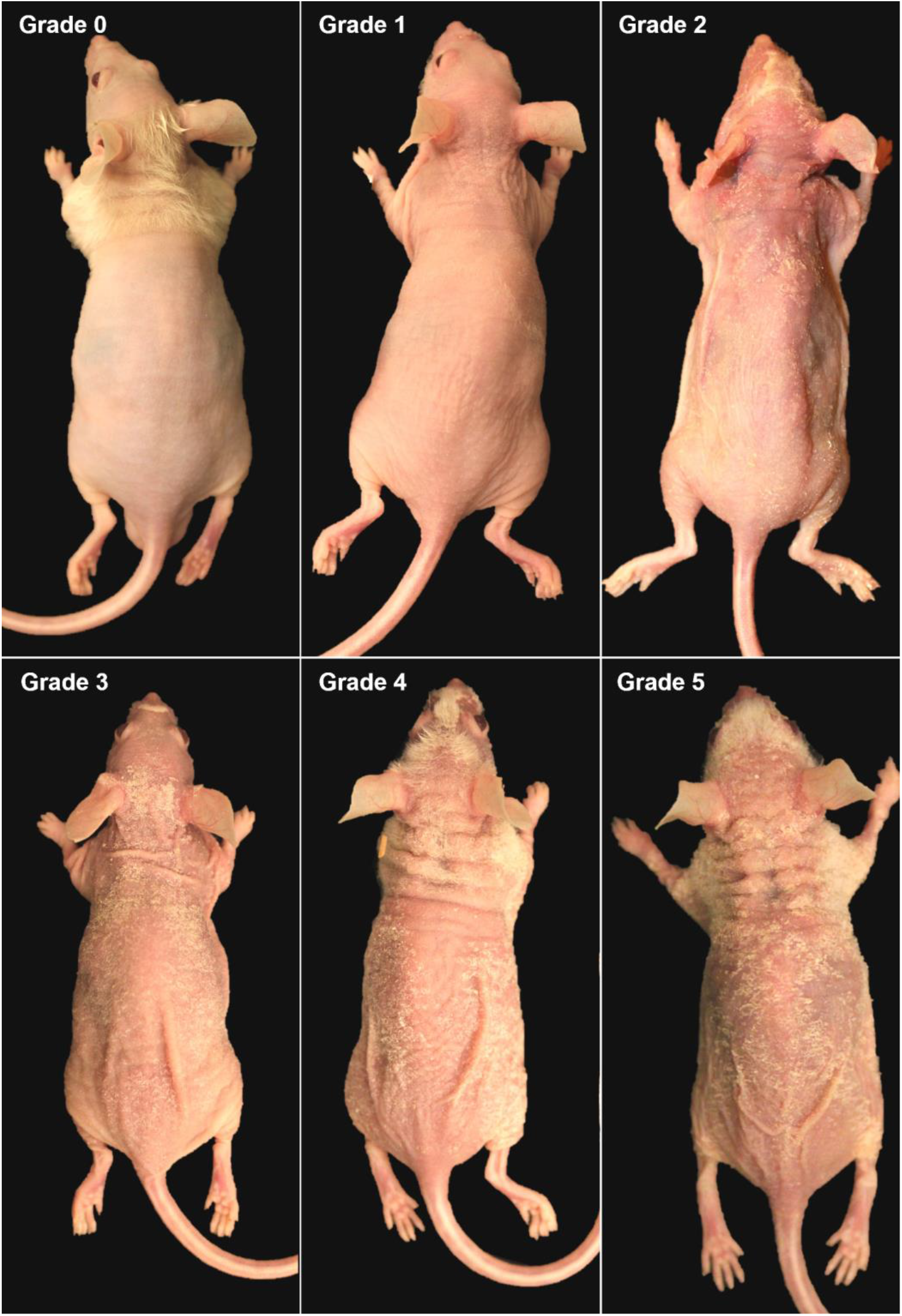
Photographs demonstrating the clinical scoring system used to assess Cb-associated skin disease in athymic nude mice at necropsy. Mice were scored 0 to 5 based on clinical signs representing mild, moderate, and severe disease. Reprinted with permission from Mendoza et al., 2023.^1^

**Figure 4:**
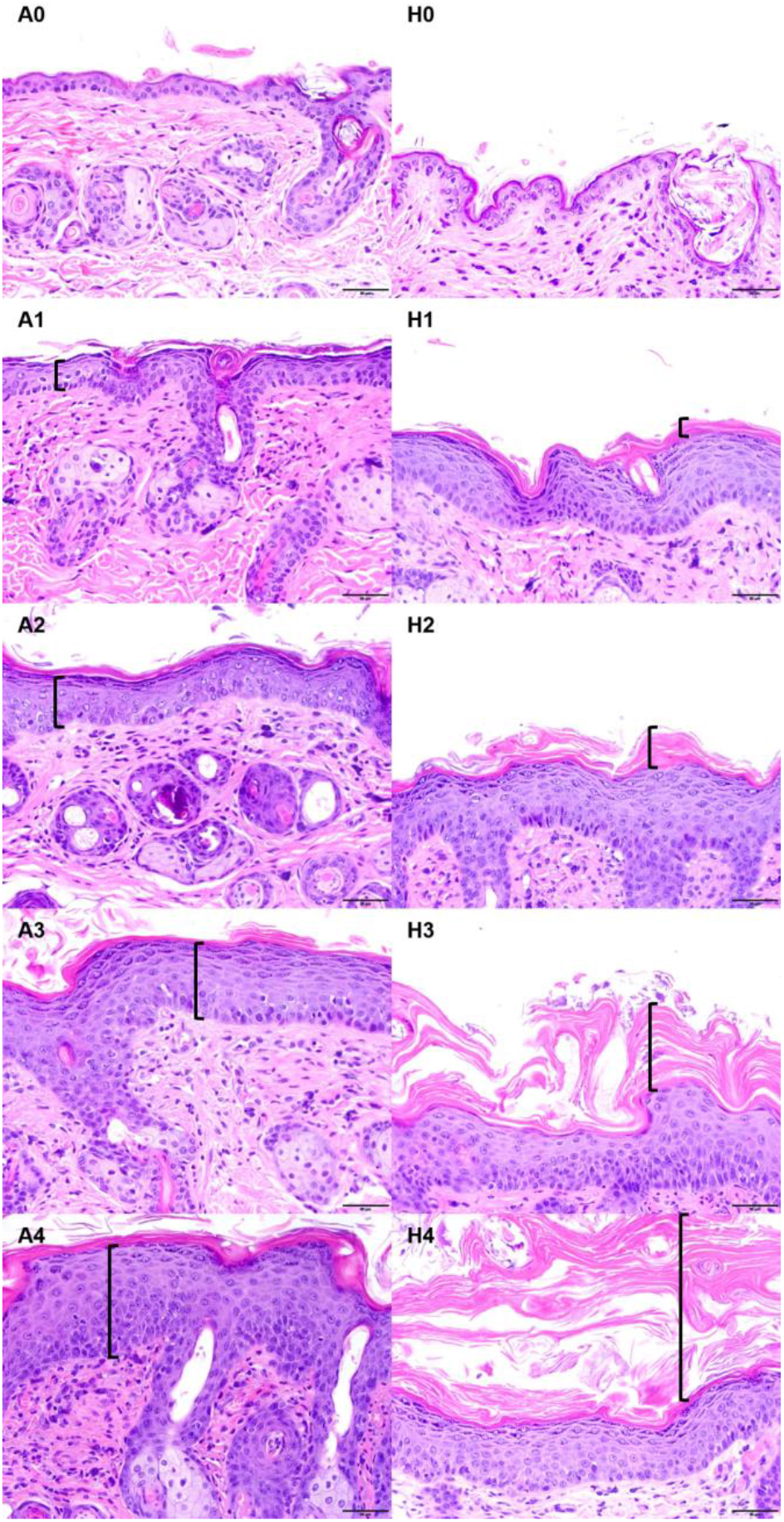
Photomicrographs demonstrating the histologic scoring system used for assessing the severity of acanthosis and hyperkeratosis in athymic nude mice infected with Cb and/or Ca. Samples were semiquantitatively scored as normal (0), minimal (1), mild (2), moderate (3), or severe (4), based on the intensity of acanthosis and orthokeratotic hyperkeratosis. A0 to A4, acanthosis grade; H0 to H4, hyperkeratosis grade; black bracket represents the location and amount of acanthosis (A) or hyperkeratosis (H). Reprinted with permission from Mendoza et al., 2023.^1^

### Postmortem gross examination and skin histopathology

On either 28 or 63 DPI with the PI, mice were euthanized by CO_2_ overdose. Each mouse underwent a complete postmortem gross examination, including macroscopic inspection of the skin and all internal organs. Full-thickness skin samples (n=4), approximately 1 cm in length, were collected from the ear, head (between the ears), and fore- and hindlimb (anterior) using a scalpel from each mouse. Specimens were fixed in 10% neutral buffered formalin, processed in alcohol and xylene, embedded in paraffin, sectioned at 5µm, and stained with hematoxylin and eosin. Histologic examination was performed on all skin biopsies to confirm the presence or absence of Cb-related lesions and to determine the magnitude of the histologic changes. Slides were evaluated and lesions recorded by a board-certified veterinary pathologist (ICM) who was blind to groups. Samples were semi-quantitatively scored as normal (0), minimal (1), mild (2), moderate (3), or severe (4), depending on the degree of acanthosis, orthokeratotic hyperkeratosis (orthokeratosis), bacteria, and inflammation with a maximum possible score of 16.^1^ Specifically, acanthosis refers to an elevated number of keratinocytes and indicates an increase in epidermal thickness resulting from hyperplasia and often hypertrophy of cells of the stratum spinosum. Minimal acanthosis was defined as an increased thickness of up to a 3-cell stratum spinosum layer; mild as 4 to 7 cell layers; moderate as 8 to 10 cell layers; and, severe as greater than 10 cell layers. Orthokeratosis refers to increased thickness of the stratum corneum without retention of keratinocyte nuclei and was mostly identified in a compact to laminated pattern. Minimal orthokeratosis was defined as an increased thickness of up to 50µm; mild as 100 to 200µm; moderate as to 200 to 300µm; and, severe as greater than 300µm. Bacterial colonies were assessed based on the presence of distinguishable clusters of short bacilli in the stratum corneum or the lumen of hair follicles. Minimal presence of bacteria was defined as small amounts of scattered bacterial rods with no formation of characteristic clusters; mild as 1 to 2 clusters; moderate as 3 to 5 clusters; and, severe as greater than 6 clusters. Inflammation was mostly comprised of a mixture of mononuclear cells and neutrophils, associated both with and without hair follicle rupture. Minimal inflammation was defined as single superficial pustules or small numbers of inflammatory cells scattered in the dermis; mild as slightly larger or increased numbers of superficial pustules or slightly greater numbers of inflammatory cells scattered in the dermis; moderate as larger or increased numbers of pustules or increased numbers of inflammatory cells in the dermis, frequently forming clusters around capillaries or adnexal units; and, severe as large and multiple pustules with high numbers of inflammatory cells forming coalescing clusters or sheets in the dermis, often infiltrating the epidermis or adnexal epithelia. For all mice, the average of each of the 4 microscopic characteristics was calculated using 4 biopsies per mouse. The “combined histopathology score” was calculated by adding the average of all 4 histologic characteristics.

### Statistical analysis

*A priori* power analysis concluded that a sample size of 5 per group would confer significance with an effect size of 3. Due to the potential for animal loss, 6 mice were utilized per group. Daily clinical scores and mean total histopathology scores, and the duration of clinical signs between NPI and Ca inoculated or NPI SB exposed groups were compared to their uninoculated, challenged and unchallenged controls. Daily clinical scores were compared between groups using Mann-Whitney U tests. The AUC was compared between groups using a one-way ANOVA and the post-hoc multiple comparisons test. Differences in duration of clinical presentation between all groups were compared using a one-way ANOVA. Differences in histologic characteristics (including hyperkeratosis, acanthosis, bacterial colonies, and inflammation) were compared between groups using Mann-Whitney U tests. The mean of histopathology scores of the 4 sections of skin for each histopathologic criteria assessed for each of the groups were also compared using Kruskal-Wallis and Dunnet’s multiple comparison tests. The PCR copy numbers of each isolate were compared across groups at each timepoint using unpaired t tests. P values <0.05 were considered statistically significant. All calculations were performed using Prism 10 for Windows (GraphPad Software version 10.3.0, San Diego, CA). Post-hoc power analysis was performed using GPower for Windows (GPower software version 3.1, Heinrich Heine University of Düsseldorf, Germany).

## Results

### Clinical score and disease progression

#### Topical Inoculation

The temporal course of clinical disease, which differed by experimental group, is shown in Figure 5. All mice receiving only the PI developed skin lesions 9 DPI reaching a MPCS of 2.5 +/- 0.5 (mean +/- SD) on 11 DPI with all but 1 mouse resolving clinically by 14 DPI. All mice from the Ca+PI group developed clinical signs later at 12 DPI with the PI reaching a lower MPCS of 1.2 +/- 0.4 on 15 DPI, with all but 1 mouse resolving clinically by 17 DPI. A single mouse from both the Ca+PI and the PI groups maintained a clinical score of 1 until the day they were submitted for histopathologic evaluation (28 DPI). In contrast, mice from the NPI+PI group never developed observable lesions. The cumulative disease burden, reflected by the area under the clinical disease curve (AUC; total area +/- SEM), was significantly higher (p<0.05) for the PI only group (AUC: 10.75 +/- 1.3) as compared to both the NPI+PI (AUC: 0) and the Ca+PI (AUC: 6.25 +/- 1.15) groups. When comparing groups that developed clinically detectable lesions (PI and Ca+PI), the former developed significantly more severe lesions earlier as the clinical signs were delayed and less severe in the latter. The duration of observable lesions (mean +/- SD) and number of mice per group presenting with clinical signs was 7.7 +/- 5.1 days; 6/6 for the mice only receiving the PI and 7.3 +/- 4.4; 6/6 for the Ca+PI group. Mice receiving sterile media, Ca only, NPI only, and NPI+PI failed to develop clinically observable skin lesions.

**Figure 5:**
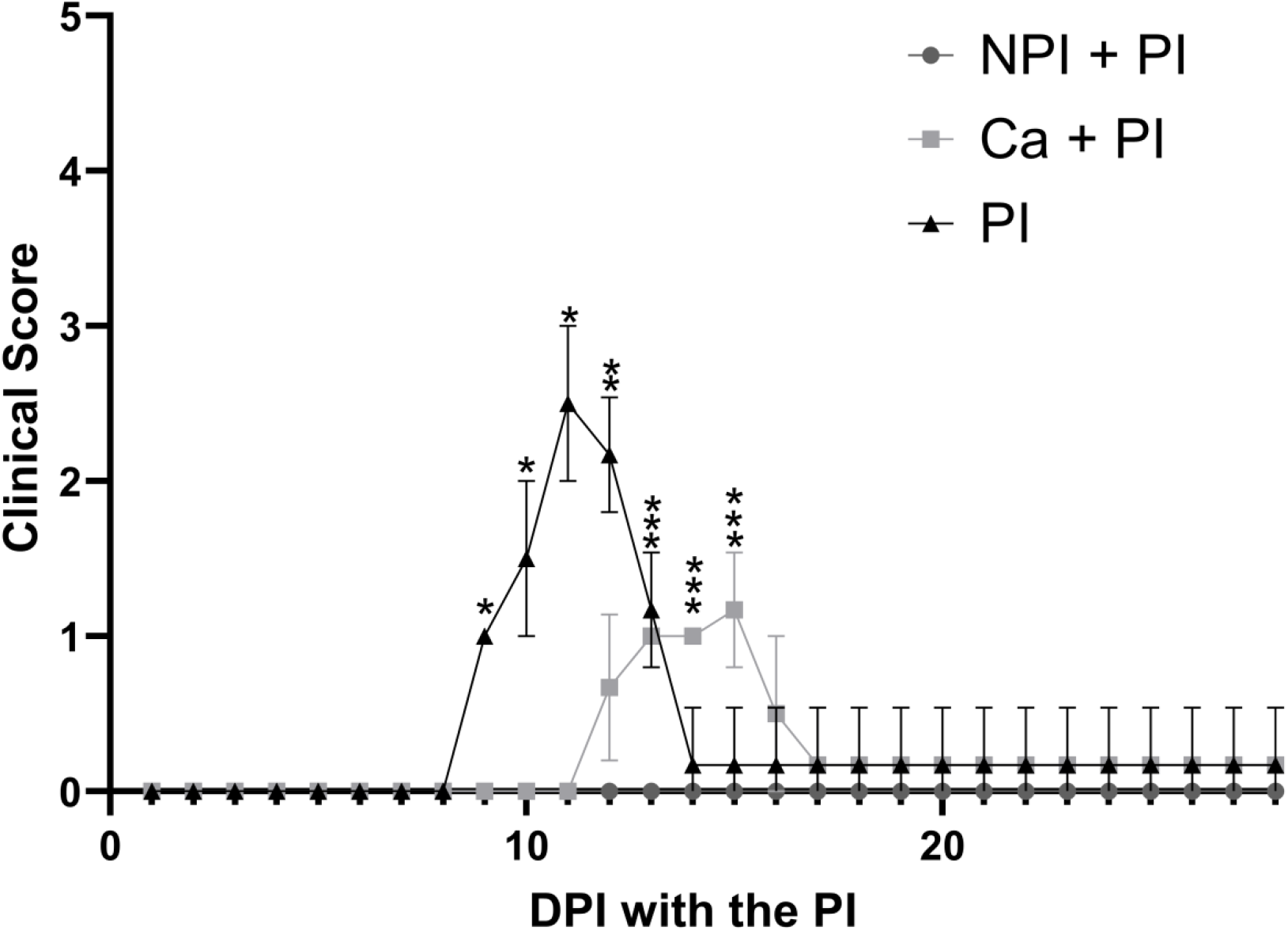
Clinical score by experimental group. Scores reflect the mean +/- SD for n=6 mice/group assessed daily. The NPI+PI and Ca+PI groups were inoculated with the PI 2 weeks after being inoculated with the NPI or Ca. *The PI clinical score was significantly (p<0.05) higher when comparing to the NPI+PI and Ca+PI groups. ** The PI clinical score was significantly (p<0.05) higher when comparing the NPI+PI and Ca+PI groups and the Ca+PI clinical score was significantly (p<0.05) higher than that of NPI+PI. *** The Ca+PI clinical score was significantly (p<0.05) higher than that of the PI and the NPI+PI groups. Mice receiving sterile media, only NPI or only Ca did not develop observable lesions and are excluded. DPI = days post inoculation; NPI = non-pathogenic Cb isolate; PI = pathogenic isolate; Ca = *Corynebacterium amycolatum*.

#### Seeded Bedding

The temporal course of clinical disease which differed by experimental group is shown in Figure 6. All mice housed on sterile bedding receiving the PI developed observable clinical lesions 9 DPI, reaching a MPCS of 2.83 +/- 0.69 on 11 DPI with observable lesions resolving by 15 DPI. None of the mice housed on NPI seeded bedding for 3 or 7 days prior to inoculation with the PI developed observable clinical lesions except for a single mouse (#SB7.5) in the SB7+PI group which developed observable skin lesions 6 DPI attaining a peak clinical score of 2, 9 DPI. Lesions resolved by 11 DPI before recurring 30 DPI before mice received the second PI inoculation. This mouse was sacrificed 35 DPI after reaching the humane endpoint with a peak clinical score of 4; the mouse also presented dehydrated with a generalized lymphadenopathy. Results (Figure 6) are presented with and without this outlier. None of the remaining mice housed on seeded bedding developed observable lesions when inoculated with the PI. Additionally, none of the mice receiving a second inoculation with the PI 35 DPI developed observable clinical lesions. The cumulative disease burden, as reflected by the AUC (AUC +/- SEM) was significantly higher (p<0.05) in mice housed on sterile bedding (9.5 +/- 1.15) as compared to mice housed on either NPI seeded bedding for 3 (0) or 7 (3.83 +/- 2.18) days, even when including the clinically affected outlier. Clinical lesions were observed in all 6 mice housed on sterile bedding and the duration of clinically observable lesions were as long as 6 days (5.33 +/- 0.52 days; mean +/- SD), whereas there were no observable lesions in mice in the SB groups except for the previously described outlier which initially presented with lesions for 5 days before developing lesions a second time for 4 days 30 DPI prior to being euthanized.

**Figure 6:**
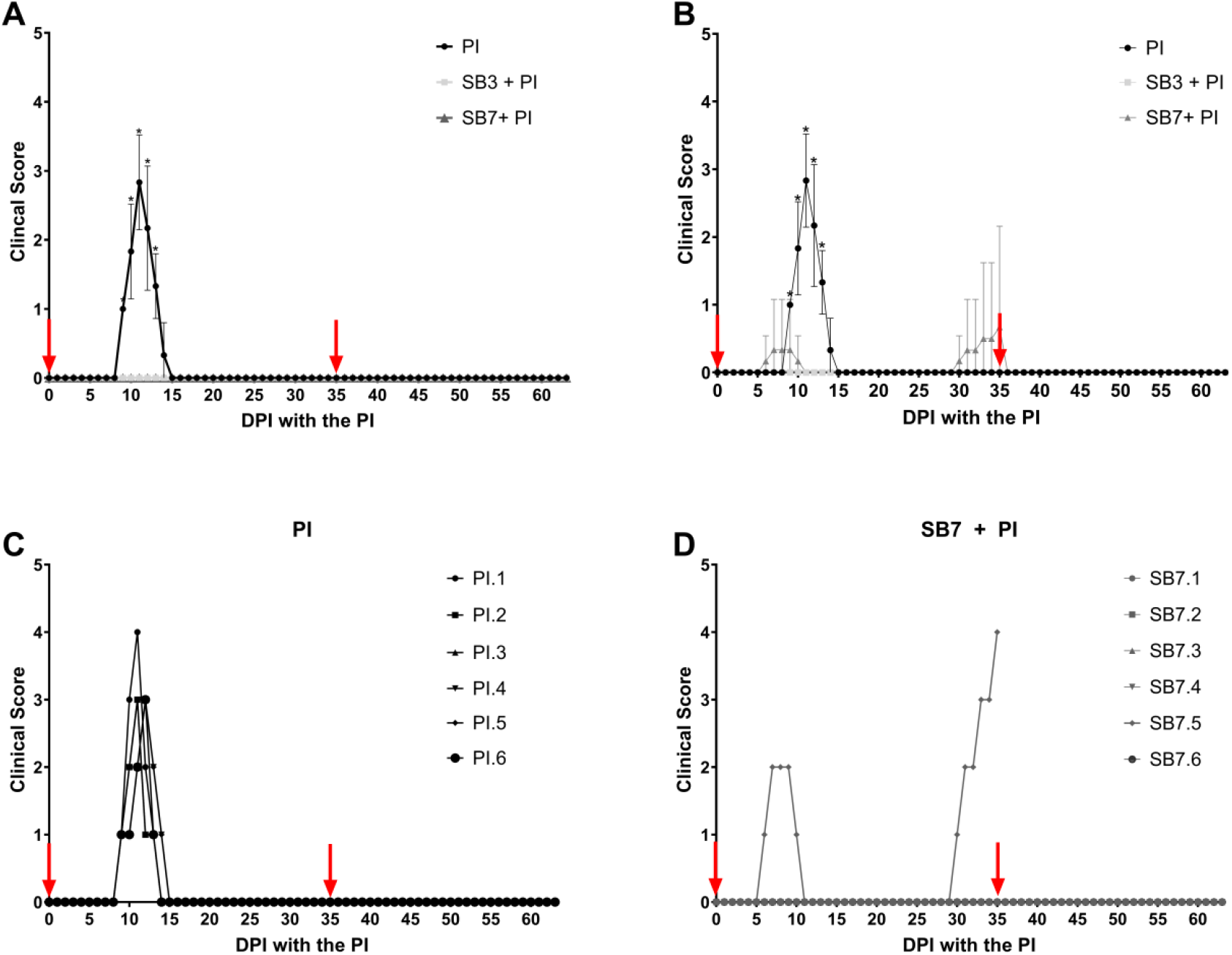
**(A)** and **(B)** Clinical score (mean +/- SD) for n=6 mice/experimental group without [A] and with [B] the outlier, respectively. Red arrows indicate inoculation days (0 and 35) with the PI. The mean clinical scores for mice housed on sterile bedding were significantly higher (*, p<0.05) than the scores of mice housed on seeded bedding after exposure to the PI on the indicated days. **(C)** Individual clinical scores for mice housed on sterile bedding following inoculation with the PI. **(D)** Individual clinical scores for mice housed on seeded bedding for 7 days prior to inoculation with the PI. A single mouse (# SB7.5) developed observable lesions 6 DPI resolving by 10 DPI only to recur with greater severity on 30 DPI. PI = Pathogenic isolate; SB = Seeded bedding; SB7 = received seeded bedding 7 days prior to inoculation with the PI; SB3 = received seeded bedding 3 days prior to inoculation with the PI.

### Bacterial culture

Semiquantitative bacterial colonization was assessed on arrival and then every 14 days following PI inoculation throughout the course of each study. All mice were negative for Cb on intake. In the topical inoculation study (Table 2), all mice inoculated with Cb were culture positive (+++) at all time points post inoculation, except for mice from the NPI+PI group 14 DPI, which had a (++) score. On 0 DPI, one cage tested (+) for Cb despite this sample being collected prior to inoculation with the PI. This result was not confirmed by PCR and given that it was found to occur in a single cage, it was presumed to reflect contamination. Ca was consistently isolated post inoculation, but the colonization varied from (+) to (+++) for mice that were also inoculated with the PI. Mice in the seeded bedding study (Table 3) were sampled 7 or 3 days after receiving the NPI seeded bedding, immediately before inoculation with the PI; mice in the SB7 group were (+++) at 0 DPI, whereas mice in the SB3 group were culture negative until 7 DPI, at which point they were (+++).

**Table 2:** Cb and Ca culture results for topical inoculation study.

| Group | -14 DPI | -7 DPI | 0 DPI | 14 DPI | 28 DPI |
| --- | --- | --- | --- | --- | --- |
| NPI | - | +++ Cb | +++ Cb | +++ Cb | NA |
| Ca | - | +++ Ca | +++ Ca | +++ Ca | NA |
| Sterile Media | - | - | - | - | NA |
| PI | - | - | +/- | +++ Cb | +++ Cb |
| NPI + PI | - | +++ Cb | +++ Cb | ++ Cb | ++ Cb / +++ Cb |
| Ca + PI | - | +Ca / +++ Ca | ++ Ca / +++ Ca | +++ Cb; +++ Ca | +++ Cb; + Ca |

All mice, except those from the PI and sterile media groups, received either Ca or NPI on -14 DPI. Mice from groups PI, NPI+PI, and Ca+PI received the PI on 0 DPI after aerobic culture sample collection. Results reflect 2 cages/group. Results per cage were the same unless reflected as x/x indicating the individual result for each cage. DPI= days post inoculation; PI=pathogenic isolate; NPI=nonpathogenic isolate; Cb=*Corynebacterium bovis*; Ca=*Corynebacterium amycolatum*; NA=not applicable as mice were sacrificed prior to the timepoint.

**Table 3:** Cb culture results for the seeded bedding study.

| Group | Intake | 0 DPI | 7 DPI | 21 DPI | 35 DPI | 49 DPI | 63 DPI |
| --- | --- | --- | --- | --- | --- | --- | --- |
| SB7+PI | - | +++ | +++ | +++ | +++ | +++ | +++ |
| SB3+PI | - | - | +++ | +++ | +++ | +++ | +++ |
| PI | - | - | +++ | +++ | +++ | +++ | +++ |
| SB | - | +++ | +++ | +++ | +++ | +++ | +++ |

Animals from groups SB7+PI, SB3+PI, and PI were inoculated with the same dose of PI on 0 and 35 DPI, after the aerobic culture sample collection. Mice from group SB7+PI received the seeded bedding 7 days prior to the initial challenge date (0 DPI), whereas mice from SB3+PI received the seeded bedding 3 days prior. DPI=days post inoculation; PI=pathogenic isolate; Cb=*Corynebacterium bovis*; SB=seeded bedding.

### PCR

#### Topical Inoculation

The copy numbers of the NPI, Ca and the PI were assessed by PCR on arrival and every 14 days until the study’s endpoint (Figure 7). Neither Cb isolate (NPI or PI) nor Ca were detected on any mouse prior to inoculation with the various bacterial strains/species used in the study. There were no significant differences in the amount of the NPI (Figure 7A) or Ca (Figure 7B) detected at 0 or 14 DPI when comparing groups of mice receiving only the initial inoculum to those which were subsequently inoculated with the PI, although there was considerable variability in the amount of the NPI detected in the NPI group 14 DPI with copy numbers ranging from 49 to 876. In contrast, there was significantly more PI present on the skin of the mice receiving only the PI 14 DPI as compared to the mice which had been inoculated with either the NPI or Ca (Figure 7C). However, at 28 DPI, the only mice which had significantly less PI present were the mice that had been inoculated with the NPI. Inoculation with Ca did not impact the amount of PI present 28 DPI.

**Figure 7:**
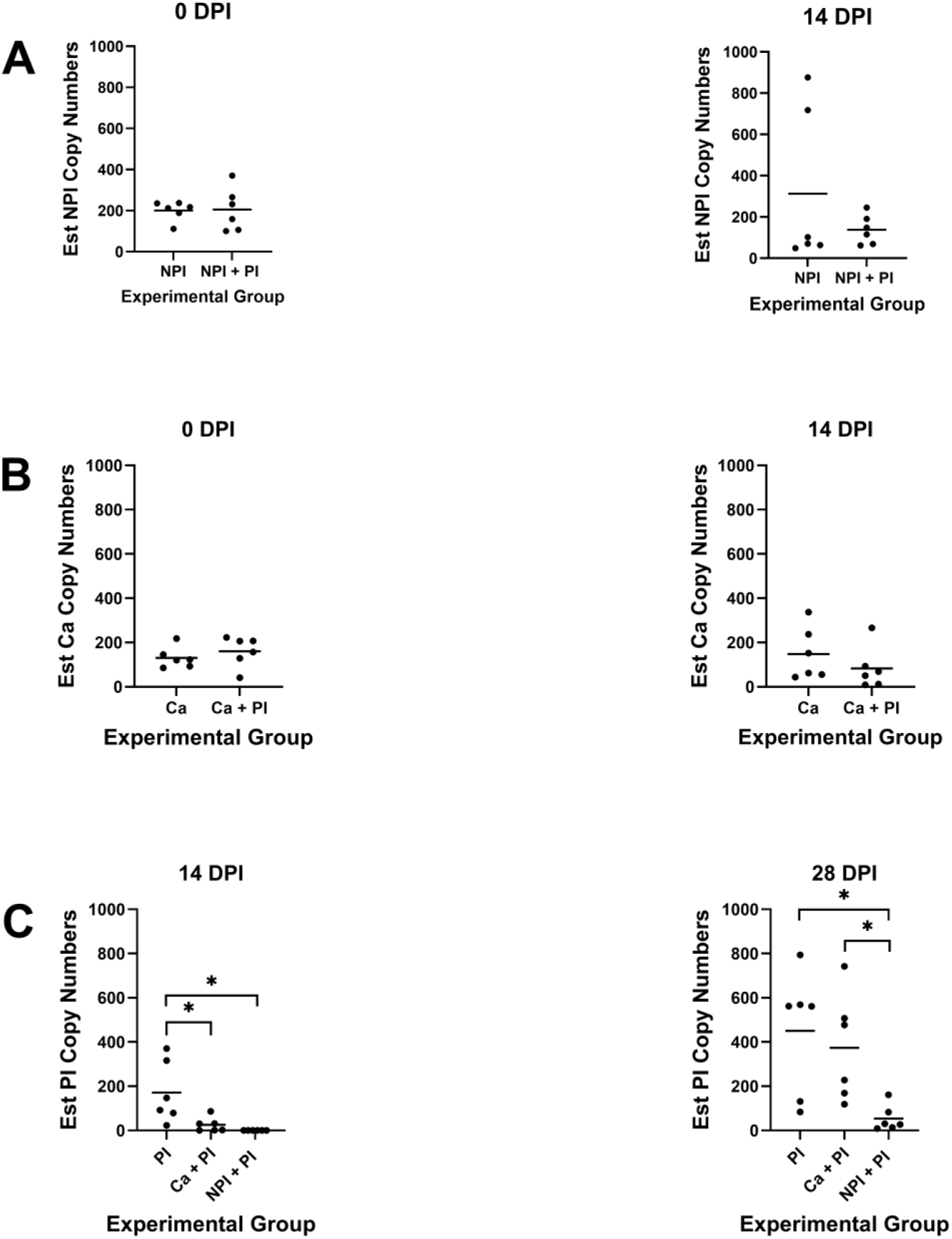
Mean estimated copy numbers of the NPI **(A)**, Ca **(B)**, or PI **(C)** at specific study time points. **(A)** There were no significant differences in NPI copy numbers in mice that received only NPI as compared to mice which were subsequently inoculated with the PI at 0 and 14 DPI. **(B)** There were no significant differences in copy numbers of Ca present on the skin of mice that only received Ca as compared to those that were subsequently inoculated with the PI at 0 and 14 DPI. **(C)** The copy numbers of PI were significantly lower in mice that had been inoculated with the NPI at 14 and 28 DPI. In contrast, Ca only impacted the PI copy numbers at 14 DPI in that they were not significantly different in mice not receiving Ca at 28 DPI. The horizontal bar represents the data mean. *, p < 0.05; DPI = days post inoculation; NPI = nonpathogenic isolate Cb; PI = pathogenic isolate; Ca = *Corynebacterium amycolatum*.

#### Seeded Bedding

The quantity of the NPI present on the mouse’s skin was evaluated by PCR on arrival as well as 3 or 7 days after receiving the NPI seeded bedding, and then every 14 days until the study’s endpoint (Figure 8). There was significantly greater amounts of the NPI on the skin of mice receiving only SB as compared to the mice challenged with the PI 3 days (SB3+PI) after receiving NPI seeded bedding at 0 and 7 DPI. Additionally, the amount of NPI in the SB group was significantly higher than both other groups (SB3+PI and SB7+PI) at 35, 49, and 63 DPI. At 21 DPI, there was significantly more NPI on the skin of mice in the SB7+PI as compared to mice in the SB3+PI group. The amount of PI detected in the PI group was significantly higher than both other groups at 7 and 21 DPI. Additionally, the amount of PI detected in mice in the SB3+PI group was significantly higher than detected in mice in the SB7+PI group at 21 DPI. The amount of NPI present on the skin of the mice decreased temporally for all mice that received it, but at a faster rate when the mice were also inoculated with the PI. The amount of PI detected was initially low or absent for mice in both SB+PI groups, but eventually the PI was detected in all groups with no significant difference between mice that did or did not receive the seeded bedding prior to exposure to the PI. Mouse #SB7.5 had similar amounts of the NPI and PI when compared to other mice in the group at every timepoint except 35 DPI, at which point there was more than 2-fold greater PI detected as compared to the mouse in the group with the next highest copy number.

**Figure 8:**
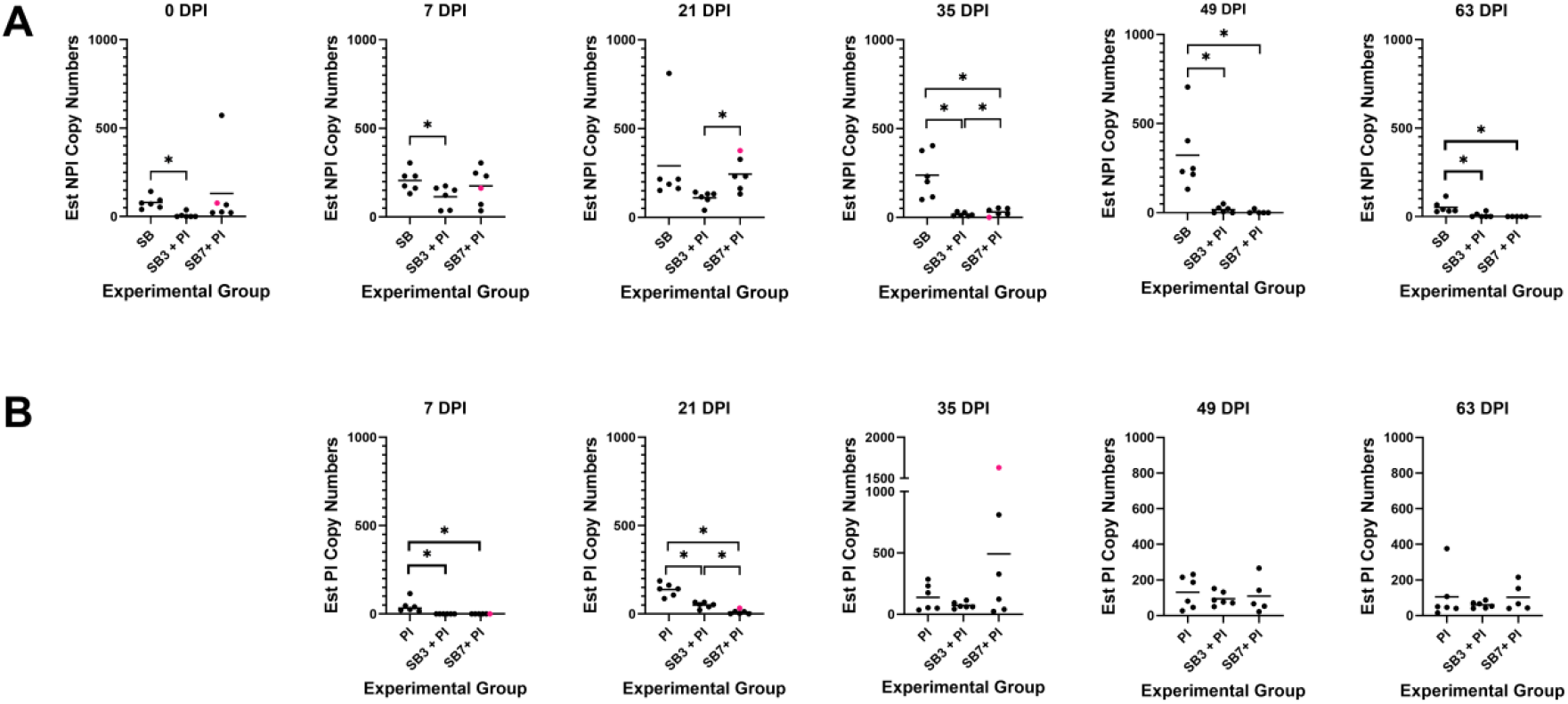
Estimated copy numbers of **(A)** the NPI isolate provided in the seeded bedding and **(B)** the pathogenic isolate (PI). The horizontal bar represents the data mean. The copy number of the outlier is reflected in pink. *, p<0.05 (statistics computed excluding the outlier) when comparing the reflected groups. DPI = days post inoculation; SB = NPI seeded bedding; PI = pathogenic isolate; SB7 = received NPI seeded bedding 7 days prior to inoculation with the PI; SB3 = received NPI seeded bedding 3 days prior to inoculation with the PI.

### Histopathology

Skin pathology was assessed 28 (topical inoculation) or 63 (seeded bedding) DPI following inoculation with the PI at which point none of the mice, except for 2 in the topical inoculation study, had clinically observable lesions. Additionally, 1 mouse from the seeded bedding study (SB7+PI) was euthanized 35 DPI as it had reached the defined humane endpoint.

#### Topical Inoculation

The histopathology scores for the topical inoculation study are provided in Figure 9. The combined histopathology scores (mean +/- SD) for the PI (7.75 +/- 0.85) and Ca+PI (7.46 +/- 1.74) groups were significantly higher than all other groups. Interestingly, the combined histopathology scores were not significantly different when comparing the PI and Ca+PI groups. The combined score for the NPI+PI (5.38 +/- 0.41) group was significantly higher when compared to the Ca (3.83 +/- 1.02) and SM (3.17 +/- 1.08) groups. The NPI group score (5.04 +/- 1.4) was significantly higher than the SM group (3.17 +/- 1.08) but there was no significant difference in the combined criteria scores between SM (3.17 +/- 1.08) and Ca (3.83 +/- 1.02) nor between NPI (5.04 +/- 1.4) and Ca (3.83 +/- 1.02) groups. When comparing individual score components, the PI and Ca+PI groups had significantly higher scores for hyperkeratosis and acanthosis as compared to Ca, NPI, and SM groups. Additionally, the NPI+PI group scores for these specific criteria were significantly higher when compared to the SM group. Bacteria were only seen in groups inoculated with Cb; therefore, the PI, Ca+PI, NPI, and NPI+PI groups had significantly higher bacterial colonies scores when compared to the Ca and SM groups. Inflammation was present in all groups with the PI group having a significantly higher score compared to the other groups except Ca+PI. The Ca+PI group had a significantly higher inflammation score as compared to the Ca and SM groups.

**Figure 9:**
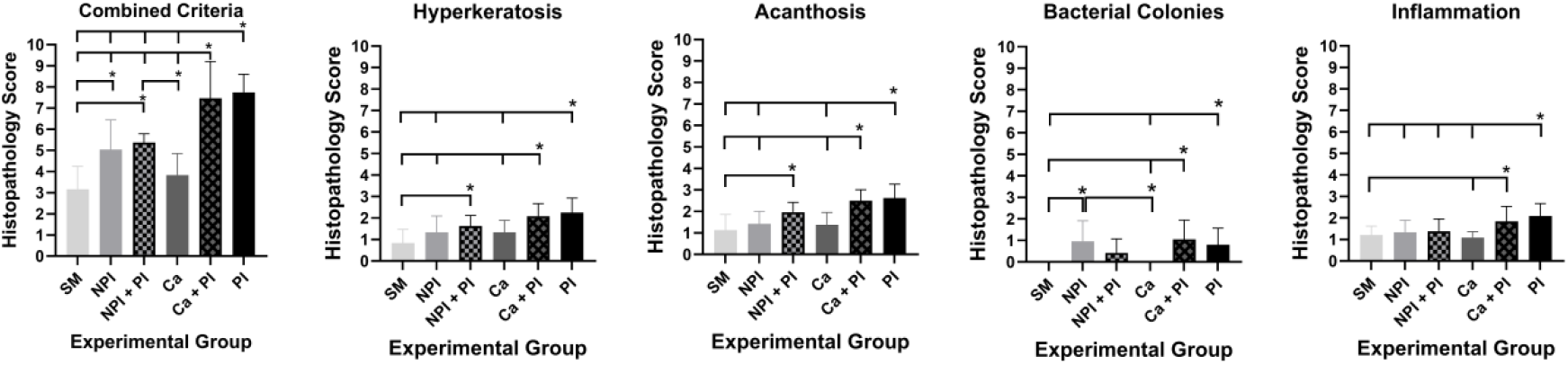
Histopathology scores (combined and individual criteria; mean +/- SD) for each group (n=6/group) in the topical inoculation study. *, p< 0.05 and is located over the group that was significantly different when compared to the other representative groups designated with tick marks on the corresponding line. SM = sterile media; NPI = nonpathogenic isolate Cb; PI = pathogenic isolate; Ca = *Corynebacterium amycolatum*.

#### Seeded Bedding

The histopathology scores for the seeded bedding study are provided in Figure 10. There were no significant differences in the combined criteria scores between groups. There was significantly greater hyperkeratosis (1.46 +/- 0.59) observed in the PI group as compared to the SB7+PI (1.05 +/- 0.51) group; the PI had significantly more severe acanthosis (1.67 +/- 0.56) as compared to the SB group (1.29 +/- 0.46). There were no other significant differences observed across groups for any specific criteria.

**Figure 10:**
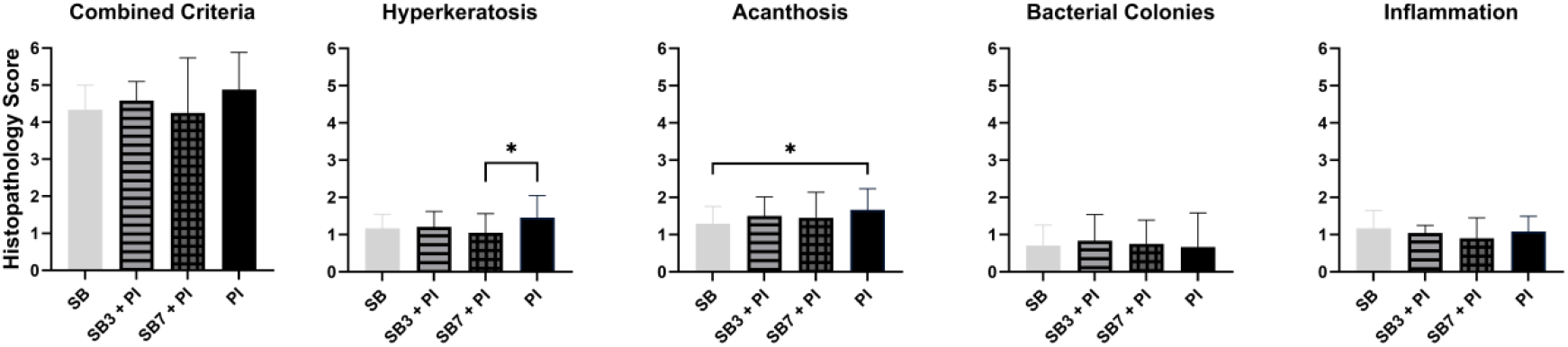
Histopathology scores (combined and individual criteria; mean +/- SD) for each group (n=6/group) in the seeded bedding study. *, p< 0.05. Mouse #SB7.5 was excluded. SB = NPI seeded bedding; PI = pathogenic isolate; SB7 = received NPI seeded bedding 7 days prior to inoculation with the PI; SB3 = received NPI seeded bedding 3 days prior to inoculation with the PI.

Mouse #SB7.5 which had been euthanized on 35 DPI, as it had reached the defined humane endpoint, had marked, multifocal, orthokeratosis and acanthosis with intracorneal bacterial colonies with colony morphology consistent with Cb, and dermal histiocytic and neutrophilic infiltrates all of which are typically observed in severe cases of CAH presentation. This mouse also had marked, multifocal lymphoid follicular hyperplasia, medullary plasmacytosis, and sinus histiocytosis indicating a generalized reactive lymphadenopathy as well as bilateral hydrocephalus.

## Discussion

This study demonstrated that prior inoculation or exposure of athymic nude mice to a nonpathogenic Cb isolate prevented clinical disease when the mice were subsequently inoculated with a pathogenic Cb isolate. It is important to recognize that although we refer to the isolates utilized in this study as NPI or PI, the isolates pathogenicity may only reflect their biological behavior in the stock of mice and their associated microbiome used in this study. The use of Ca was similarly evaluated as we had observed that mice from a vendor’s breeding colony in which Ca is a component of the skin microbiota failed to develop CAH when the colony became inadvertently infected with Cb. Additionally, we recently found that nude mice from a colony in which Ca is a component of the microbiota failed to develop CAH when infected with the PI, while the same nude mouse stock maintained at a different geographical site free of Ca developed CAH when infected with the same dose of the PI.^5^ While Ca showed promise in that the bacterium provided some degree of protection, the NPI was far superior in preventing the development of clinical signs. Therefore, we subsequently evaluated the NPI to protect against subsequent PI challenge by seeding bedding with the bacterium as a “proof of principle” practical method for implementing our finding.

Topically inoculating mice with the NPI provided complete protection against clinically observable CAH when mice were subsequently challenged with the PI, as no mice from this group demonstrated clinical Cb-associated disease. While the quantity of the NPI measured by PCR was not significantly different when comparing PI challenged to unchallenged mice at the time points evaluated, there was significantly less PI found on the skin of mice previously inoculated with the NPI at 14 and 28 DPI suggesting that the NPI inhibits PI colonization and/or growth, as well as the associated disease. Aerobic culture was of minimal value as it cannot distinguish between the Cb isolates utilized in this study; however, it did reflect the durability of Cb colonization.

Even though mice inoculated with both the NPI and PI did not demonstrate clinical disease, they did have mild Cb-associated histopathologic skin changes as their composite score was significantly higher than those in mice in the negative control group inoculated with sterile media. However, in terms of individual components, only NPI’s score for bacterial colonies was significantly higher than those in the negative control group; the NPI’s scores for the other components: hyperkeratosis, acanthosis, and inflammation were not significantly different from those of the negative control. Additionally, their composite and select individual scores were significantly lower than mice in the positive control group, suggesting that the NPI was effective in limiting the PI from causing greater skin pathology. Importantly, as the mice in the positive control group were sacrificed after their skin disease resolved or improved markedly, had mice in both groups been evaluated and compared at earlier time points, the differences in skin pathology would likely have been considerably more dramatic.

In contrast, while inoculating mice with Ca delayed the onset and decreased the severity of clinical CAH when compared to mice in the positive control group receiving only the PI, it was not as effective as the NPI in preventing disease. The amount of PI detected by qPCR in mice inoculated with both bacterial species was not significantly different than mice inoculated with only the PI at either time point evaluated, indicating that the PI did not impact the colonization with Ca. However, while Ca impeded colonization and growth of the PI initially when assessed at 14 DPI, 2 weeks later the amount of PI detected by qPCR had rebounded and was not significantly different than the PI inoculated positive control. This finding was consistent with the temporal appearance of clinical disease, which was delayed in mice infected with both Ca and the PI. Whether the lower, but not statistically different, amount of PI found on the skin of Ca and PI infected mice at 28 DPI was enough to reduce the clinical disease score could not be ascertained but is a reasonable assumption.

Mice inoculated with only Ca had minimal skin histopathology as their composite and individual scores, except for the presence of bacteria, were not significantly different as compared to mice receiving only sterile media. However, mice inoculated with Ca and the PI had composite and select individual pathology scores that were significantly higher than all the other groups, except for mice in the PI inoculated positive control group, thus Ca does not prevent Cb-associated histopathologic changes. The delay and decrease in clinical presentation does suggest that Ca’s presence can provide some clinical benefit; however, when the qPCR and histopathology data are taken into consideration, Ca did not offer a sufficient degree of protection to warrant further study and we were not able to recapitulate our observation that mice from colonies in which Ca was a component of the skin microbiota did not develop CAH when inoculated with the PI.^5^ However, it is unclear if it is Ca alone or whether Ca serves as a marker for other skin microbiota that alone or collectively with Ca mitigates the impact of Cb.

When considering a practical method of inoculating nude mice with the NPI we could utilize in our nude mouse colony, we sought a method that would be easy to replicate and implement. In addition to seeding bedding, we considered inoculating the enrichment material we provide. Bedding was selected as we found it easier to disperse the inoculum in bedding and thought it more likely that all cage occupants would be exposed to the NPI as we have observed that not all mice interact with the enrichment material we employ. Additionally, Manuel et al. successfully infected athymic nude mice by transferring soiled bedding from mice with clinically apparent CAH and we routinely detect Cb in sentinel mice housed on soiled bedding when evaluated in our colony health monitoring program.^6^ An additional benefit is large batches of seeded bedding could be created and stored at -80°C for extended periods (data not shown). We selected 2 intervals post-seeded bedding exposure to challenge mice with the PI. The intervals evaluated, 3 and 7 days, reflect the earliest and most likely intervals after arrival that nude mice would be exposed to enzootic Cb circulating in our colony. The shorter 3-day interval reflected the earliest interval post-arrival that nude mice should be handled by investigative staff as we recommend a 72h post-arrival acclimatization period; the longer 7-day interval reflects our cage change frequency.

The NPI-seeded bedding (SB) led to colonization of all groups of mice with the isolate; however, there were varying amounts of NPI detected among groups temporally. In general, unchallenged mice had greater NPI detected throughout the experimental period. Interestingly, the amount of NPI detected by qPCR decreased as early as 35 DPI in the PI inoculated mice and, at the last time point (63 DPI) the amount of NPI detected in the unchallenged group decreased to levels approximating that measured in the PI challenged groups, although the difference, albeit small, remained significant.

With respect to the detection of the PI, more PI was detected in unchallenged mice as compared to those inoculated with the isolate at earlier post-exposure time points. However, over time, the differences among groups became insignificant. Importantly, despite finding decreasing amounts of NPI on the skin, none of the mice exposed to the PI a second time at 35 DPI developed clinical signs. This finding demonstrated the robustness of the protection provided by the initial NPI exposure and/or its initial impact was sufficient to allow the host to develop protective responses. We have only observed recurrence of CAH experimentally in Cb-infected nude mice when the mice were repetitively administered the immunosuppressant, cyclophosphamide.^2^

The NPI seeded bedding provided near complete protection against clinical CAH, as only a single mouse developed Cb-associated clinical disease. Importantly, this mouse (#SB7.5) was ultimately euthanized as it developed severe skin lesions, generalized lymphadenopathy, dehydration, and lethargy 35 DPI which likely accounted for its’ idiosyncratic CAH. The quantity of PI measured on this mouse at euthanasia was significantly higher than other mice in this group and, conversely, NPI was not detected on the mouse’s skin. These findings also correlated with the mouse’s peak clinical score. This mouse was found to have hydrocephalus, which is uncommon in athymic nude mice, as it is most commonly documented in B6 mice and to the authors’ knowledge has not been documented to occur spontaneously in nude mice.^24^ It is interesting to speculate that the hydrocephalus could have resulted from systemic inflammation associated with Cb infection with the associated inflammation causing post-infectious hydrocephalus resulting from obstructed CSF flow.^25^ However, this is not a documented outcome in nude mice infected with Cb, so it is likely that the unique presentation was multifactorial, and the mouse’s immune system was likely further compromised, contributing to the atypical presentation.

In contrast to the topical inoculation study in which the cumulative pathology scores were significantly lower in NPI + PI inoculated mice as compared to mice receiving only the PI, there were minimal discernable differences in composite scores among groups in the seeded bedding study. One reason may reflect the longer duration of the latter study in which mice were euthanized at 63 DPI versus those in the former at 28 DPI. The longer post inoculation interval with the PI may have led to a further decrease in pathologic severity.

Irrespective of the method by which the NPI was administered, we have shown for the first time that Cb-associated clinical signs can be mitigated by exposing mice to an NPI prior to inoculation with a PI. Importantly, there were limitations of this study. A single investigator (AM) scored the clinical lesions and was not blinded to the groups. However, the clinical scoring system was well defined. Additionally, only one purported pathogenic Cb, nonpathogenic Cb or Ca isolate was evaluated in a single nude mouse stock with its associated microbiome. Other bacterial isolates or nude mouse stocks/microbiomes could have generated disparate results. How the NPI provides protection requires further study. Whether it protects through competitive exclusion and/or an immunologic response remains plausible, however an immune response is more likely. The principle of competitive exclusion postulates that one species will eventually outcompete the other. However in the seeded bedding study, the magnitude of and decrease in colonization of the NPI observed over time did not correlate with the level of protection the isolate provided, suggesting competitive exclusion is less likely, although the initial competition between the NPI and PI may have provided sufficient time for the mouse to mount an immune response obtunding the clinical signs and pathology typically seen following infection with the PI.

In humans, keratinocytes can detect harmful bacteria and induce production of antimicrobial peptides (AMP), like RNase 7, to control the growth of pathogenic bacteria on the skin.^18,19^ While RNase 7 expression is currently restricted to primates, it is possible there is a murine equivalent AMP that could serve a similar function.^26^ Similarly, a murine homologue of human beta-defensin, an epithelial protein with antimicrobial properties, has been found to be inducible upon introduction of a pathogen.^27^ While specific mechanism(s) by which the NPI provides protection are unknown, it is possible that the NPI may trigger one or more of these immunologic pathways, contributing to the clinical protection. Another consideration implicating the role of the immune system is our finding in a pilot study that the NPI did not protect highly immunodeficient NSG (NOD.Cg-*Prkdc*^*scid*^ *Il2rg*^*tm1Wjl*^/SzJ) mice from Cb-associated disease and pathology when inoculated with the PI.

Implementing the study’s findings requires further inquiry. It is unclear as to whether these experimental observations will translate to a nude mouse colony in which the mice may be further immunosuppressed because of experimental manipulations, and different and perhaps more virulent Cb isolates may be circulating. A clinical trial would be the likely next step. Given the practicality and feasibility of seeded bedding, we would utilize this method of exposure. However, even if the NPI was shown to be effective in a trial, concerns remain about the impact of introducing a bacterium that can induce skin pathology and perhaps cause other effects that may confound research. Additionally, the isolate could potentially revert to become pathogenic and problematic.

## Abbreviations

AUC: Area under the curve
Cb: *Corynebacterium bovis*
Ca: *Corynebacterium amycolatum*
CAH: *Corynebacterium*-associated hyperkeratosis
DPI: Days-post-inoculation
CFU: Colony forming unit
NPI: Nonpathogenic isolate
MPCS: Mean peak clinical score
PI: Pathogenic isolate
SB: Seeded bedding
SD: Standard deviation
SEM: Standard error of the mean

## Acknowledgements

We would like to acknowledge John D’Allara, Sotirios Dayanis, Kate Fodor, Glory Leung, Noah Mishkin, Michael Palillo, Amanda Ritter, and the staff from the Laboratory of Comparative Pathology for their assistance with various technical components of the study. Additionally, earlier conversations with staff at The Jackson Laboratory (Mert Aydin, Anthony Mourino, and James Fahey) regarding the microbiome of their colony, which includes Ca and their experience with a nonpathogenic Cb isolate provided important background for this study.

